# Remote Deep Brain Stimulation by Transgene-free Magnetomechanical Approach

**DOI:** 10.1101/2021.11.27.470141

**Authors:** Chih-Lun Su, Ping-Hsiang Yen, Chao-Chun Cheng, Po-Han Chiang

## Abstract

Various physical stimulation methods are developed to minimize the invasiveness of deep brain stimulation (DBS)^1–3^. Among them, only magnetic field can penetrate into the biological tissues without scattering or absorption^4^, which makes it ideal for untethered DBS. Recently developed magnetogenetics have shown the potential of developing treatments for neurological disorders^5^. However, magnetogenetic approaches have potential side effects from overexpression of exogenous ion channels and gene delivery with viral vectors^6,7^. Here, we demonstrated that the iron oxide magnetic nanodiscs (~270 nm) can be used as transducers to trigger calcium responses in the wild-type cultured neurons during the application of slow varying weak magnetic fields (50 mT at 10 Hz). Moreover, we identified that the intrinsic mechanosensitive ion channel transient receptor potential canonical (TRPC), which were widely expressed in the brain^8^, plays the main roles in this magnetomechanical stimulation. Finally, when we applied magnetic fields to the awake mice with magnetic nanodiscs injecting into subthalamic nucleus, the magnetomechanical stimulation triggered neuronal activities in the targeted region and the downstream region. Overall, this research demonstrated a magnetomechanical approach that can be used for wireless neuronal stimulation *in vitro* and untethered DBS in awake mice *in vivo* without implants or genetic manipulation.

## Main Text

Conventional electrical deep brain stimulation (DBS) has been used for treating neurological disorders, especially motor disorders like Parkinson’s diseases, essential tremor and other disease^9^. However, the using of electrical stimulation requires invasive chronic implantations with electrode into the deep brain regions^10^. To minimize the invasiveness of DBS, accumulating approaches, including optical^1^, acoustic^2^ and electromagnetic^3^ neuronal modulation approaches, were developed. In optogenetics, the lights for activating opsins can be scattered and absorbed easily by biological tissues. The implantation of optical fiber is necessary to deliver lights into deep tissues. In acoustic approaches, like sonogenetics and focus ultrasound stimulation, the ultrasound waves can be scattered, reflected and distorted by skulls and bones. Mounting of ultrasound probe with aqueous cranial window are required in the acoustic neuronal stimulation. Only magnetic fields can penetrate into the brain without absorption or scattering^4^. Transcranial magnetic stimulation (TMS) is a non-invasive neuronal stimulation approaches by using strong magnetic fields (>1 T) to induce electric currents in the brain. But clinical TMS is limited to cortical stimulation. The strong magnetic fields used by TMS cause undesirable side effects like muscle twitch, facial pain and other discomforts^11^. In the last decade, magnetic approaches with weak magnetic fields by using nanoparticle-based neuronal modulation were developed rapidly^5,12–16^.

Magnetothermal stimulations use the heat dissipated from magnetic nanoparticles via hysteretic power loss with application of alternative magnetic fields at radio frequency (100 kHz to 1 MHz)^13^. To manipulate the neuronal activity with magnetothermal stimulations, thermosensitive ion channels, transient receptor potential vanilloid 1 (TRPV1) or anoctamin1, were overexpressed in the target neurons^5,13,14^. Lately, another magnetic approach, magnetomechanical stimulation, was demonstrated in both peripheral nervous system (PNS) and central nervous system (CNS). Mechanosensitive ion channel, Piezo1/2 and TRPV4, is highly expressed in sensory neurons in PNS^17^. Previous study showed that by using ~250 nm magnetic nanodiscs with weak and slow varying magnetic field (<25 mT at 5 Hz), the torque of magnetic nanodiscs could induce Ca^2+^ responses in mechanosensitive neurons in primary dorsal root ganglia (DRG)^15^. In contrast to PNS, Piezo1/2 expression in CNS neurons is very low. By overexpression Piezo1 in the brain, neurons could be stimulatesd by the torque of 500 nm magnetic nanoparticles with 20 mT magnetic field at 0.5 Hz^16^. However, the potential side effects of overexpression exogeneous gene are still unclear. The viral vectors for gene delivery also raise safety concerns in application^6,7^. Therefore, in this study, we developed a non-genetic approach to eliminate the necessity of gene delivery.

The transient receptor potential canonical (TRPC) is a non-selective cation channel family. There are 3 subfamilies: TRPC1/4/5, TRPC2, and TRPC3/6/7. They are involved in neuron developments, learning, memory and fear related behaviors^8,18^. Amount them, mammalian TRPC1, 5 and 6 are mechanosensitive and play a role in stretch-stimulated responses^19–21^. Moreover, TRPC are widely expressed in various brain regions^8,22^. In compare with Piezo1/2, the TRPC requires higher mechanical force^23^. In this study, we hypothesize that with slightly stronger magnetic field intensity than that used for magnetomechanical stimulating Piezo1 in previous studies^15,16^, we could induce TRPC-mediated neuronal responses (Fig. 1A) for neuronal modulation *in vitro* and *in vivo.* To transduce mechanical force to the neurons, we used previously described iron oxide magnetic vortex nanodiscs (MNDs)^15^. Which had better colloidal stability in absence of magnetic fields. In weak magnetic fields at low frequency, the MNDs could be used as transducers to activate the mechanosensitive ion channels on the cell membrane^15^.

**Figure 1.**
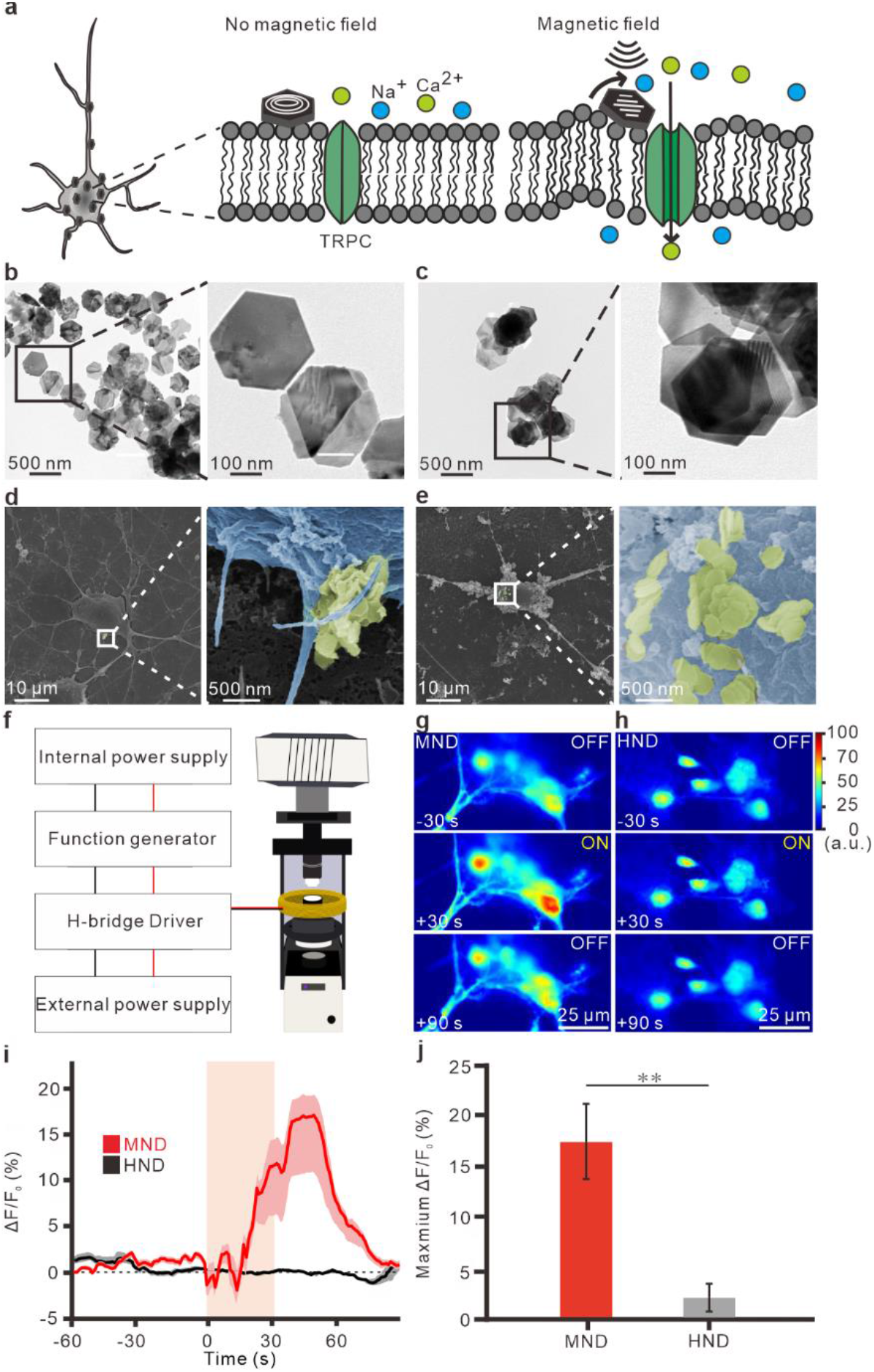
Magnetomechanical induced neuronal activity with MNDs. **a**, Schematic of magnetomechanical stimulation by using MNDs. **b** and **c,** TEM images of PMAO-coated MNDs (b) or HNDs (c). **d** and **e**, SEM images of MNDs (d) and HNDs (e) on the membrane of cultured neurons. **f**, Schematic of magnetic apparatus for fluorescence microscope. **g** and **h**, Color maps of fluorescence intensity of neurons with MNDs (g) or HNDs (h) application. Magnetic field with 50 mT at 10 Hz was applied for 30 s. **i**, Average traces of Ca^2+^ responses by magnetic field stimulation. Red line, MND group. Black line, HND group. Light area, s.e.m. **j**, Maximum change of Ca^2+^ responses in different groups. **p<0.01.

MNDs were synthesized by a two steps synthesis protocol^15^ (Fig. S1). The hematite nanodiscs (HNDs) was first synthesized with 180 °C in autoclave reactor. Next, the MNDs were produced by reduction of HNDs (Fig. S1B). With different volume of water during synthesis, we were able to synthesis different size of MNDs (Fig. 1B and S1H-I). To generate larger mechanical force, we were using nanodiscs with larger diameters (~270 nm) for magnetomechanical stimulation (Fig. 1B-C, S1H). To functionalize the nanodiscs, all the nanodiscs used for neuronal stimulation, including MNDs and HNDs, were coated with poly(maleic anhydride-alt-1-octadecene) (PMAO)^15^. The negative charged nanoparticles could facilitate the attachment of nanodiscs to the excitable neuron cells^24^. After PMAO coating, the zeta potential of the coated nanodiscs were −53.5 ± 3.7 mV for MNDs and −54.5 ± 2.3 mV for HNDs (Fig. S1J). When we applied the nanodiscs (70 μg/ml) on the primary hippocampal neurons. Both MNDs and HNDs attached to the membrane of primary hippocampal neurons (Fig. 1D-E). A custom-designed magnetic apparatus for fluorescence microscope was used for magnetic stimulation under upright microscope (Fig. 1F, S2). The neuronal activities were measured by using Ca^2+^ indicator, Fluo-4. We found that when we apply MNDs to the cultured neurons, a magnetic stimulation with 50 mT at 10 Hz induced Ca^2+^ responses in neurons (Fig. 1G, I-J). In contrast, when we applied HNDs to the neurons, the magnetic field with same condition cannot induce any Ca^2+^ responses (Fig. 1H-J).

To identify the optimal condition for magnetomechanical stimulation with intrinsic mechanosensors, we compared the MND-mediated neuronal responses by the magnetic fields with different frequencies and field intensities (Fig. 2). The alternative magnetic fields from 1 to 20 Hz were used to induce Ca^2+^ responses in cultured neurons with MND (70 μg/ml). At different frequency, the magnetic field intensities were sequentially increased from 10 mT to 50 mT (Fig. 2A-D, S3). We found that the Ca^2+^ responses with 50 mT simulation was significantly larger than other magnetic field intensities from 10 to 40 mT (Fig. 2E). The Ca^2+^ responses at 5 to 20 Hz were significantly larger than responses at 1 Hz (Fig. 2E). When stimulating neurons at 50 mT with different intensities (Fig. 2F-I), we found that 10 and 20 Hz stimulation could induce larger Ca^2+^ responses than 1 and 5 Hz stimulation (Fig. 2J, S4A). 10 Hz stimulation could activate more cell population than other conditions (Fig. 2K, S4B). In contrast, HNDs cannot induce any responses in all conditions (Fig. 2F-J, S4C-D). These results indicated that 50 mT at 10 Hz is the optimal condition to activate neurons with magnetomechanical stimulation. Moreover, when we applied the magnetic stimulation repeatedly for 4 times, we observed multiple Ca^2+^ responses in cultured neurons (Fig. 2F-I, S4). After repeated magnetic stimulation for 4 times, the cell viability were tested with propidium iodide, a small fluorescent molecule that only can penetrate into the cell membrane of dead cells but not live cells^25^. We didn’t observe any cell death after stimulation at different frequencies in either MNDs or HNDs treated neurons. (Fig. S5).

**Figure 2.**
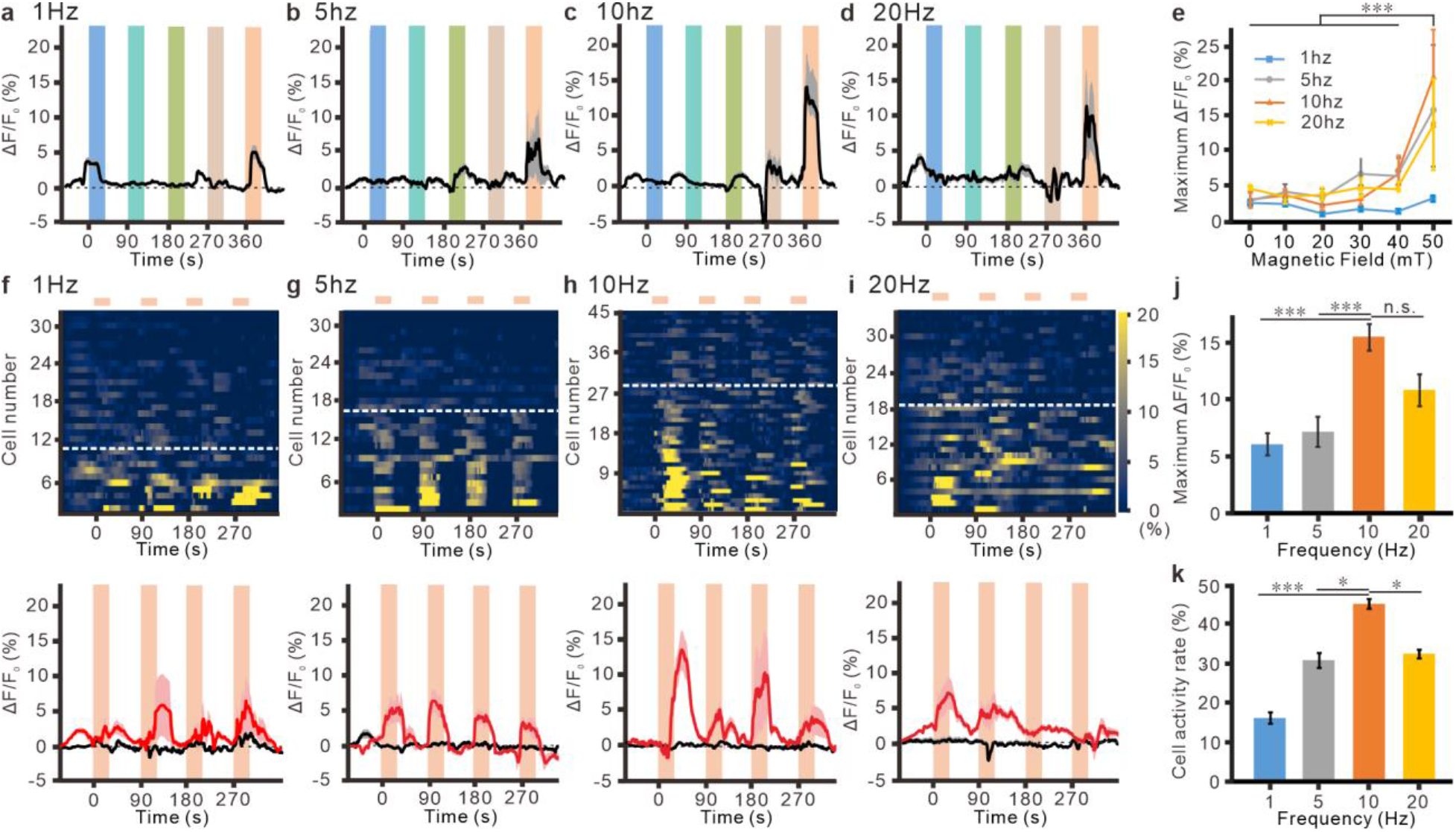
Tuning the magnetic field intensity and frequency for wireless stimulations. **a** to **d**, The averaged traces of MND-mediated neuronal Ca^2+^ responses which were stimulated by different frequency at 1 Hz (a), 5 Hz (b), 10 Hz (c), or 20 Hz (d). In each frequency, magnetic field intensity was sequentially increased from 10 to 50 mT. The gray areas are s.e.m. **e**, The maximum ΔF/F0 at different conditions. F = 9.639, p < 0.001 for field intensity, F = 6.155, p < 0.001 for frequencies; F = 1.459, p = 0.11 for interaction of frequencies and intensities. Two-Way ANOVA. **f** to **i**, Multiple stimulations with 50 mT at 1 Hz (f), 5 Hz (g), 10 Hz (h), or 20 Hz (i). Top, Heatmap of Ca^2+^ responses in individual neurons. The cells below white dash lines were activated during stimulations. Bottom, The averaged traces of fluorescence change. Red line, MND group. Black line, HND group. Light area, s.e.m. **j**, The maximum change of fluorescence at different frequencies. p < 0.001, Kruskal-Wallis test. **k**, The cell activity rate at different frequencies. p < 0.001, Kruskal-Wallis test. *p < 0.05, ***p < 0.001.

There are several mechanosensing cation channels are expressed in the mammalian cells. including Piezo1/2, TRPC, TRPV4, ASIC3^26,27^. Among them, TRPC and TRPV4 are reported in the CNS ^26^. In contrast, Piezo1/2 and ASIC3 are mainly expressed in the PNS^27^. To investigate the mechanism of the MND-mediated responses, pharmacological approach was used to dissect the ion channels that involved in the magnetic stimulated responses in hippocampal neurons. First, with Ca^2+^-free extracellular solution, all the magnetomechanical induced Ca^2+^ responses are abolished (Fig. 3A, S6). It indicated that these MND-mediated responses were contributed by Ca^2+^ influx from external solution. In TRPC family, TRPC1, 5 and 6 are reported as mechanosensitive cation channel^19,20^. We found that a specific blocker for TPRC family, SKF-96365 (50 μM), eliminated the magnetomechanical induced Ca^2+^ responses (Fig. 3B, S6). In addition, other non-specific TRPC blockers were also used to exam the contribution of TRPC, including GsMTx4, d-GsMTx4 (5 μM), and 2-Aminoethoxydiphenyl borate (2-APB). GsMTx4 and d-GsMTx4 are antagonist for Piezo1 and Piezo2, respectively. They are also antagonists for TRPC1,5 and 6^21^. 2-APB is TRPC antagonist and TRPV1-3 agonist, but insensitive to TRPV4^28^. Similar to SKF-96365, all the TRPC blockers, including GsMTx4 (5 μM), d-GsMTx4 (5 μM) and 2-APB (100 μM), were able to eliminate the magnetomechanical induced responses in hippocampal neurons (Fig. S7). These results indicated that intrinsic TRPC in hippocampal neurons plays critical roles in magnetomechanical stimulations (Fig. 3G).

**Figure 3.**
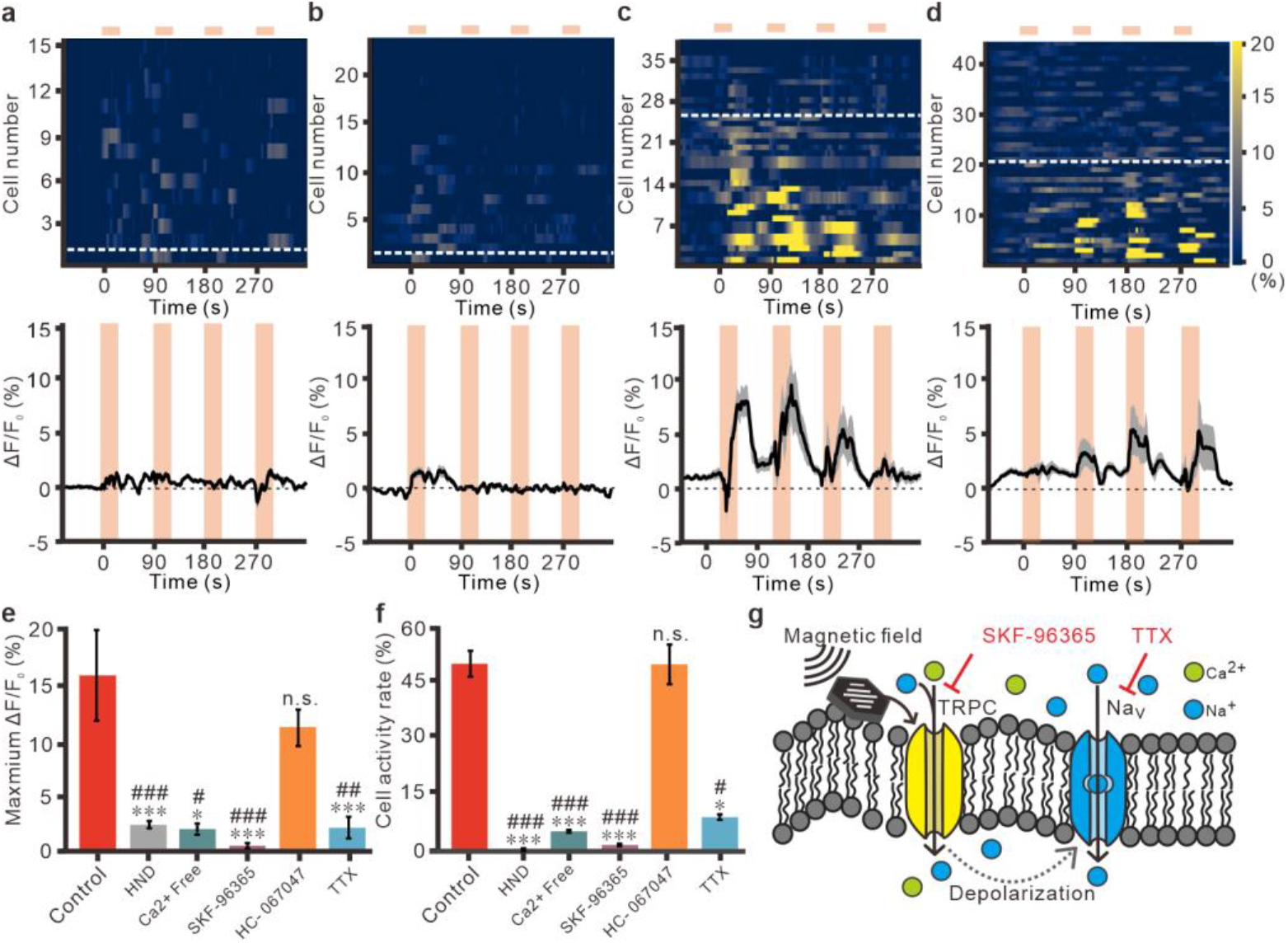
Pharmacological dissection of magnetomechanical stimulated responses in neurons. **a** to **d**, The Ca^2+^ responses by multiple magnetomechanical stimulation with bath application of Ca^2+^-free solution (a), SKF-96365 at 50 μM (b), HC-067047 at 1 μM (c) or TTX at 100 nM (d). The varying magnetic fields are 50 mT at 10 Hz for 30 s. Inter-stimulation intervals are 60 s. Top, heatmap of individual cells responses. Bottom, averaged fluorescence changes. The gray areas are s.e.m. **e**, The maximum fluorescence changes at the first magnetic stimulation. p < 0.001, Kruskal-Wallis test. **f**, The cell activity rate at the first magnetic stimulation. p < 0.001, Kruskal-Wallis test. **g**, Schematic for the mechanism of MND mediated responses. *p < 0.05, ***p < 0.001, compared to control group; #p < 0.05, ##p < 0.01, ###p < 0.001, compared to HC-067047 group.

Next, the role of TRPV4 in magnetomechanical stimulation in hippocampal neurons were investigated by using TRPV4 specific blocker, HC-067047^15^. However, HC-067047 (1 μM) cannot modulate the MND-mediated responses (Fig. 3C, S6). Which indicated that the magnetic stimulated responses were not mediated by TRPV4. Finally, the voltage-gated sodium channel blocker, tetrodotoxin (TTX; 100 nM), was used for investigating whether action potentials were involved in the magnetomechanical induced Ca^2+^ responses. The first stimulation induced Ca^2+^ responses and cell activities are almost eliminated with TTX application (Fig. 3D). This result indicated that the activation of TRPC by MNDs depolarized the membrane to induce action potentials in neurons. However, when we applied magnetic stimulations for multiple times, the maximum fluorescence changes or cell activity rates were significantly reduced but not completely abolished by TTX application (Fig. S6). TRPC is a non-selective cation channel which permeable to Ca^2+^. This result indicated that we might recruit more TRPC with multiple stimulations. Overall, from the pharmacological study, we found that the torque of MND generated by alternative magnetic field can induce Ca^2+^ influx from external solution and can induce action potentials in hippocampal neurons. This magnetomechanical stimulated responses were mainly mediated by intrinsic TRPC (Fig. 3G).

Subthalamic nucleus (STN) is the target of conventional DBS with electrical stimulation for treating patients with Parkinson’s diseases^29^. Previous study showed that the mechanosensitive TRPC are largely expressed in STN^30^. To demonstrate the magnetomechanical neuronal modulation for DBS *in vivo,* nanodiscs (2 μl of 1 mg/ml) were unilaterally injected into STN of mice (Fig. 4A). After 5 to 7 days, the nanodiscs injected mice were placed into a large custom-made round coil with 20 cm inner-diameter and 25 cm height (Fig. 4B, S8). The awake mice were stimulated by magnetic field with 50mT, 10Hz with 30 sec on-30 sec off cycle for 10 min (Fig. 4B bottom). We found that the immediate early gene, c-fos, expressions in MNDs injected STN was significantly larger than contralateral STN (Fig. 4C-E). In contrast, there were no difference of c-fos expressions between the ipsilateral and contralateral STN of HNDs injected mice (Fig. 4F, S9A-B). The ipsilateral/ contralateral ratios of c-fos expressions in STN of MNDs injected mice were also significantly more than HNDs injected group (Fig. 4G). Entopeduncular nucleus (EP) is one of the downstream of STN glutamatergic projecting neurons in mouse. Which is homologous to internal Globus Pallidus (GPi) in human. Similar to STN, c-fos expressions in the ipsilateral EP of MNDs injected mice were significantly more than contralateral EP (Fig. 4H, S9C-D). But in HNDs injected mice, there were no increase of c-fos expression in EP (Fig. 4I, S9E-F). The ipsilateral/ contralateral ratios of c-fos expressions in EP of MND injected mice were also significantly more than HNDs injected group (Fig. 4J). These results showed that by using magnetomechanical approach with MNDs, we were able to wirelessly modulate the neuronal circuit in the deep brain region *in vivo*.

**Figure 4.**
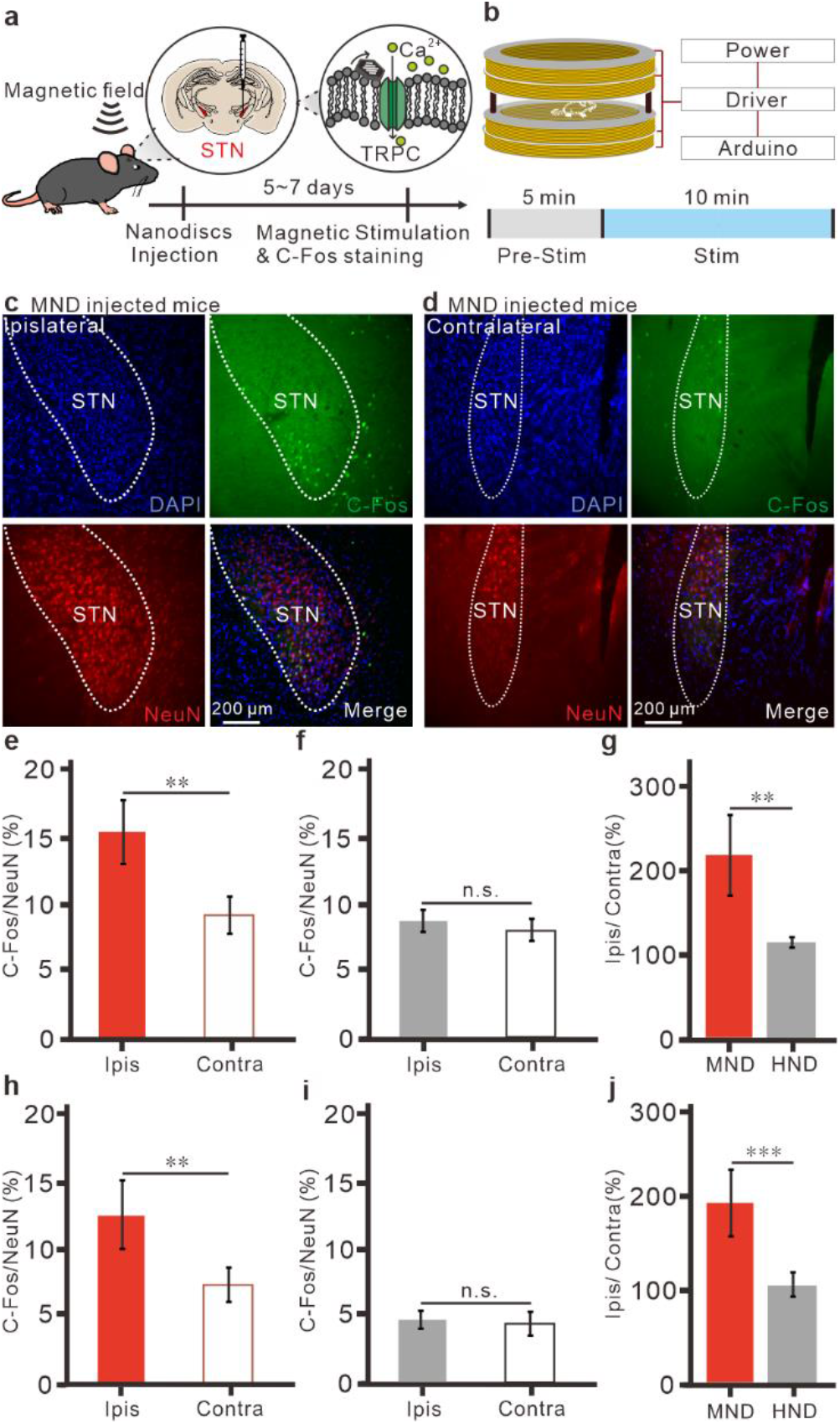
magnetomechanical stimulation *in vivo.* **a**, Schematic of magnetomechanical stimulation at STN *in vivo.* Nanodiscs are unilaterally injected into STN. Bottom, Timeline of stereotaxic injection, magnetic stimulation and c-fos staining **b**, Schematic of wireless magnetic stimulation apparatus for magnetomechanical stimulation *in vivo*. Bottom, Timeline for magnetic stimulation. **c** to **d**, Immunostaining of c-fos (top-left), NeuN (top-right), DAPI (bottom-left) and merged image (bottom-right) in STN after magnetic stimulations. Ipsilateral STN of MND injected mice (c). Contralateral STN of MND injected mice (d). **e**, c-fos/NeuN of STN in MND injected mice. **f**, c-fos/NeuN of STN in HND injected mice. **g**, The difference of c-fos/NeuN between ipsilateral and contralateral STN in nanodiscs injected mice. **h**, c-fos/NeuN of EP in MND injected mice. **i**, c-fos/NeuN of EP in HND injected mice. **j**, The difference of c-fos/NeuN between ipsilateral and contralateral EP in nanodiscs injected mice. **p<0.01, ***p<0.001, n.s., no significant.

In conclusion, we found that when applying weak and slow alternative magnetic fields (50mT at 10Hz), the torque of magnetic nanodiscs induced the activity of wild-type neurons. These magnetomechanical responses were mainly mediated by the intrinsic mechanosensitive cation channel TRPC. Finally, the activities of deep brain regions were increased by MND-mediated magnetomechanical stimulation in awake mice *in vivo*. The low intensity magnetic field varying at low frequency can easily penetrate to the deep brain region for neuronal modulation^4^. The magnetic apparatus for magnetomechanical approach is scalable to larger volume. The custom-made coil for *in vitro* and *in vivo* experiments in this study have 3.5 cm and 20 cm inner diameter, respectively (Fig. S2, S8). This scalable feature is ideal for future applications in larger animal models or humans. Iron oxide magnetic nanoparticles are clinically approved as contrast agents in magnetic resonance imaging (MRI)^4^. The iron oxide magnetic nanodiscs in this study have similar chemistries to those clinically approved nanoparticles. Therefore, our findings provide a transgene-free and minimal-invasive untethered DBS approach that can be used for understanding of neuronal circuitry in animal models and for developing treatments for neurological diseases.

Recent studies of magnetomechanical stimulation in Peizo 1-expressing DRG or overexpression Piezo1 in CNS only requires <23 mT at 1~5 Hz. In this study, we didn’t observe obvious responses with magnetic field at < 40 mT (Fig. 2A-E). The magnetic field intensity for inducing MND-mediated TRPC activity in our study was larger (50 mT) than previous researches. Which is in line with previous report that TRPC requires stronger mechanical force than Piezo1^23^. Although TRPC can response to mechanical stimulation^31^. It is debatable that whether TRPC is the direct sensor for the mechanical stimulations or not^32^. We cannot rule out the possibility that TRPC might be activated by mechanical stimulation indirectly when we apply magnetomechanical stimulation with MNDs. Nevertheless, our results revealed that the neuronal activities induced by magnetomechanical stimulation more than 50 mT were depends on the intrinsic TRPC. Although TRPC is widely expressed in the brain, if we want to target different brain regions to develop treatments for various neurological diseases, the differences of TRPC expression levels in different regions and cell types must be considered.

## Acknowledgements

We thank P. Ankieeva, D. Gregurec, and F. Koehler for materials and electronics advices.

## Author contributions

Conceptualization: PHC, CLS. Methodology: PHC, CLS, PHY. Investigation: CLS, PHY, CCC. Formal analysis: CLS. Visualization: CLS, PHY. Funding acquisition: PHC. Project administration: PHC. Supervision: PHC. Writing – original draft: PHC. Writing – review & editing: PHC, CLS, PHY, CCC.

## Data and materials availability

All data are available in the main text or the supplementary materials.

