## Supplementary Materials for "Remote Deep Brain Stimulation by Transgene-free Magnetomechanical Approach"

**This PDF file includes:**

Materials and Methods
Figs. S1 to S9

### Materials and Methods

Hematite and Magnetite Nanodiscs Synthesis

Nanodiscs synthesis process follow the protocol described in previous study[15]. The synthesis process consists two steps. First, hematite nanodiscs were synthesized by mixing 10 ml 99.5% ethanol, 0.6 ml dd$H_2O$ (or 0.8 ml dd$H_2O$ for smaller nanodiscs), 0.8 g of anhydrous sodium acetate (Sigma-Aldrich) and 0.273 g of $FeCl_3 \cdot 6H_2O$ (Sigma-Aldrich). After the mixture was homogenized by stirring, the mixture was transferred and sealed into a Teflon-lined steel vessel. The vessel was heated in the oven at 180 °C for 18 h. The hematite nanodiscs were washed with dd$H_2O$ twice. Then nanodiscs were washed with ethanol twice. Nanodiscs were dried in a vacuum desiccator. Hematite nanodiscs was further reduced into magnetite nanodiscs or used for control experiments. For reduction, 1 mg hematite nanodiscs adding 20 ml of tri-octylamine (Acros Organics) and 1 g of oleic acid (Sigma-Aldrich). The mixture was placed in a three-neck flask connected to a Schlenk line to heated to 370 °C for 25 min in an atmosphere of $H_2$ (5% with 95% Argon, Chiah Lung) and $N_2$ (99.9%, Chiah Lung). During reduction, the red hematite nanodiscs turned to dark gray. When discs cooled, the discs washed with hexane (Alfa Aesar). The magnetite nanodiscs were then dispersed in chloroform (J.T. Baker) and stored in a glass vial at 4 ºC

PMAO-coating and water transfer

First dissolved 10 mg 30,000 mw PMAO (419117, Sigma-Aldrich) in 1 mL chloroform. Then the solution is added to dry 1mg nanodiscs. This mixture sonicated for 1hr until the nanodiscs were well suspended, magnetite dissolving in PMAO to dry in vacuum desiccator overnight. Added TAE (Tris-acetate-EDTA, Biomate) buffer 4 mL with 1mg magnetite particles and sonicate at 80 ºC for 3 hr, then pelleted by using 10 min microcentrifuge and afterwards wash twice in water.

Nanodiscs characterization

Nanodiscs soaked in chloroform were dried in a vacuum desiccator and then dissolved in dd$H_2O$. The aqueous nanodiscs were placed on a copper grid (Ted Pella Inc.) and used transmission electron microscopy (TEM) for visualizing the morphology of the discs. TEM was performed with HT7800, Hitachi, Japan by microscopy core laboratory at Chang Gung Memorial Hospital (CGMH). Diameter and thickness of nanodiscs were measured by ImageJ from TEM image, each hexagonal nanodiscs were drawn three diagonals and chose longest distance as one nanodisc diameter measurement. 3.5 µl of 20 mg/ml nanodiscs were loaded on hippocampal neurons at 7 days in vitro (DIV) for 15 minutes to attach nanodiscs on cell membrane. Then use 1× PBS (Gibco) to wash three times and fixation buffer made by CGMH (3% glutaraldehyde, 2% paraformaldehyde in 0.1 M cacodylate buffer) for fixation. Fixed sample need to store in 4 ºC freezer until using field emission scanning electron microscopy (FE-SEM) to record nanodiscs attaching on neurons. FE-SEM was performed with SU8220, Hitachi, Japan by microscopy core laboratory at CGMH. FE-SEM was used to record nanodiscs attaching on neurons at 2,000× magnification and the surface features of the nanodiscs for 30,000× magnification. Nanodiscs with PMAO-coating dissolved in dd$H_2O$ were used zeta potential for detecting potential property. Zeta potential was measured by electrophoretic light scattering with DelsaTM Nano C Particle Analyzer (Beckman Coulter).

Primary hippocampal neuronal culture

All animal experimental procedures were approved by the Institutional Animal Care and Use Committee (IACUC) of National Yang Ming Chiao Tung University (NYCU). All pregnant

Sprague-Dawley rats were from LASCO. The pups of Sprague-Dawley rats at postnatal <1 days were used for primary hippocampal culture. Before seeding, 12 mm round glass coverslips were coated with 75 µl of matrigel solution, which prepared from 30x diluted matrigel (354234, Corning) with neurobasal w/o B27 supplement (157504001, Gibco), and were incubated at 37 °C for > 1 hr. Hippocampus were extracted in cold dissection solution (160 mM NaCl, 5 mM KCl, 1 mM $MgSO_4$, 4 mM $CaCl_2$, 5 mM HEPES, 5.5 mM glucose, pH was adjusted to 7.4 by NaOH). After extraction, tissues were transferred and incubated at 37 °C in prewarmed digestion solution (1 mM L-Cysteine, 0.5 mM EDTA, 1 mM CaCl2, 1.5 mM NaOH, and 10 units/ml papain (76220, Sigma-Aldrich) for 25 min. Digestion was inactivated by removing the digestion solution and incubating the tissues in inactivation solutions ( 0.25% bovine albumin and 0.25% trypsin inhibitor, 0.4% D-glucose, and 5% fetal bovine serum (6140079, Gibco) in Minimum Essential Medium (MEM w/Earle's salts w/o L-glutamine; 11090-081, Gibco) at 37 °C for 2 min. After removing the inactivation solution, tissues were triturated in serum medium (0.4% D-glucose, and 5% fetal bovine serum in MEM w/Earle's salts w/o L-glutamine) with fire polished glass pipetted (111096, Kimble). The dissociated cells were filtered by cell-drainer (93070, SPL) and incubated in the serum medium at 37 °C before seeding. After counting, cells were seeded on the materigel coated coverslips in 24-well plates with ~110000 cells/well. Hippocampal cells were cultured in neurobasal medium with B27 supplement and GlutaMAX (35050-061, Gibco). On 3 days in vitro (DIV) 20µL mitotic inhibitor (5-fluoro-2'-deoxyuridine (Sigma); 4 µM in neurobasal medium) was added to inhibit glial cells. All the imaging and stimulation are performed at DIV 5-14.

Calcium imaging with nanodiscs

Tyrode's solution (125 mM NaCl, 2 mM KCl, 2 mM $MgCl_2$, 2 mM $CaCl_2$, 25 mM HEPES, 51 mM D-glucose (Sigma-Aldrich)) was prepared for all calcium imaging for cultured cells. Fluo-4 Calcium Imaging Kit (Invitrogen) was used for measuring the calcium responses of cultured neurons. The protocol for fluo-4 imaging was indicated by Invitrogen. Fluo-4 was diluted 1000× in Tyrode's solution (Fluo-4 final concentration at 1 mM) for neurons incubate in this solution for 15-30 min, and then transfer to Tyrode's solution without Fluo-4 for imaging. 3.5 µl magnetite nanodiscs or hematite nanodiscs at a concentration of a 20 mg/mL was added to each well with 496.5µL solution in 24- wells plate. The final concentration of nanodiscs is 70 mg/well. After 5min incubation, the coverslip with hippocampal neuron coverslip was transferred to custom stage for applying magnetic field under fluorescence microscope (SS-1000-00, Scientifica). Thorlabs light source (LEDD1B), Hamamatus C13440 camera, GFP filter cube (39002, Chroma) was used for fluo-4 imaging.

Calcium Imaging Analysis

Calcium activity video were collected on an upright fluorescence microscope (SS-1000-00, Scientifica) in the ".avi" format using the HCImage software (Hamamatsu). Video's frame rate was 1 Hz. The videos were processed with a custom python script based on opencv2. The cells are autoselected by the following process: "Blur", "AdaptiveThreshold", "Erosion", "Dilation" and "findContours". The mean value of intensity in individual cells are measured and output as ".txt" file. A custom python script based on numpy was used for converting fluorescence intensity into $\Delta F/F_0$ by adapting the algorithm described previously (Gregurec et al., 2020). Where $\tau_0 = 3$ s, $\tau_1 = 5$ s, and $\tau_2 = 30$ s. The activated cell was defined by the cell with maximum $\Delta F/F_0$ more than 10 % at indicated time periods.

Pharmacology Experiments

The Tyrode's solution without calcium (125 mM NaCl, 2 mM KCl, 2 mM $MgCl_2$, 2.5 mM EGTA, 25 mM HEPES, 51 mM D-glucose (Sigma-Aldrich)) was prepared for calcium free experiments. The TRPC family inhibitor SKF-96365 (Tocris) was added to Tyrode's solution with calcium at a concentration of 50 μM. The TRPV4 specific inhibitor HC-067047 (Sigma) was added to Tyrode's solution at a concentration of 1 μM. The Tetrodotoxin (TTX) sodium channel blocker (Abcam) was added to Tyrode's solution at a concentration of 1 nM. The TRPC 1/6 and piezo1 specific inhibitor GsMTx4 (Abcam) was added to Tyrode's solution at a concentration of 5 μM. The TRPC 1/6 and piezo2 specific inhibitor D-GsMTx4 (Tocris) was added to Tyrode's solution at a concentration of 5 μM. The TRP inhibitor 2-APB (Tocris) was added to Tyrode's solution at a concentration of 100 μM. In pharmacology experiments, cultured cells were transferred from Tyrode's solution with Fluo-4 to abovementioned solutions for more than 5 min before imaging.

Animal experiments and Stereotaxic injection

All the animal experiments were approved by NYCU IACUC, in accordance with the Guide for the Care and Use of Laboratory Animals of NYCU. All C57BL/6 male mice were from LASCO, and animals were maintained under a 12 h light-dark cycle at NYCU Laboratory Animal Center before experiment. Stereotaxic injection and Animal experiment also completed in the animal center. PMAO-coated magnetite and hematite nanodiscs at 1 mg/ml were used for stereotaxic injection. Unilateral injected into STN(AP: -1.5, ML: -2.06, DV: -4.5).In experiment we injected with mice with 2 μl magnetite and hematite using an air pressure injection pump (Kd Scientific) and Hamilton microinjection syringe (7803-05, Hamilton). 4-7 days later, mice were placed in a custom-made coil to for magnetic stimulation *in vivo*. Mice were sacrificed 90 min after stimulation.

Fixed brain slicing and immunostaining

Brains were collected following transcardial perfusion of 1X PBS and 4% PFA. 50 μm coronal brain sections were cut with a vibratome (5100 MZ, Campden) with amplitude 0.5, frequency 50 hz. The brain slices were washed with PBS for three times, the slices were permeabilized with 2% (v/v) Triton X-100 (Sigma-Aldrich) for 15 min. Background was cleared up with 2% Triton-X-100, 30 % $H_2O_2$ and methanol for 10 min; After washed the slices with PBS for three times, the slices were blocked by 3% normal goat serum in PBS for 90 min at room temperature. After washed the slices with PBS for three times, slices were incubated with 1$^{st}$ antibody solution with 1:750 rabbit anti-c-Fos monoclonal antibody (9F6#2250, Cell signaling), 1:150 mouse anti-NeuN antibody (clone A60, #MAB377, MERCK), 1 % normal goat serum and 2% Triton-X 100 in PBS. Sliced were incubated at 4°C for 16-18 h. After three times washes of the cells with PBS, slices were incubated with matching secondary antibody in the PBS (1:500 goats anti-rabbit Alexa Fluor 488 (ab150113, Abcam) and 1:500 goat anti-mouse Alexa Fluor 594 (ab150116, Abcam)). After washed the slices with PBS for three times, slices were mounted on glass microscope slides by mounting solution with DAPI (GTX30920, Genetex). Leica DMI3000 inverted microscope, LED light source (pE300, CoolLED), Hamamatus C13440 camera, filter cubes (39000, 19008, 31002, Chroma) was used for imaging.

Magnetic apparatus for fluorescence microscope

The coil was an air-core coil made by 2000 turns of 18AWG self-bonding cooper wire (SBWR, Chientai). The inner-diameter, outer-diameter and height of the coil were 3.5cm, 13.5 cm and 4 cm, respectively (Fig. S2A). The coil had 7Ω Resistance and 60mH inductance. At 1 Hz varying magnetic field stimulation, 0.6A to 2.8A was used for 10 to 50 mT stimulation. At 5 Hz magnetic stimulation, 0.6A to 2.9A was used for 10 to 50 mT stimulation. At 10 Hz magnetic stimulation, 0.6A to 3A was used for 10 to 50 mT stimulation. At 20 Hz magnetic stimulation, 0.7A to 3.2A was used for 10 to 50 mT stimulation. Temperature change less than 1 ℃ in 2 minutes with 3A DC current. (Fig S2D). A custom-made H-bridge driver was used for generating varying magnetic fields with coil. The H-bridge is made up by two p-channel MOSFET (IRF4905, International Rectifier) and two n-channel MOSFET (IRF3710, International Rectifier). Beside these four MOSFET, two more n-channel MOSFET were used to control P-channel MOSFETs that in H-bridge. Range of working voltage is control by two pairs of resistance and parameter of MOSFET. Circuit can easily be modified by replacing different resistance or MOSFET. Function generator (33210A, Keysight) was used for generating ±5 V square waves. The signal from function generator passed through a voltage follower and an inverter (TLC2272, Texas Instrument). The gains of voltage follower and inverter equal to one. The phases of signals from voltage follower and inverter have 180° difference. Each signal controls a half bridge of the full-bridge driver. Internal power supply (PS-3030DF model, LONGWEI) was used for op amps on the custom-made H-bridge driver. External power supply (IT6721, ITECH) was used for generating magnetic field in the coil.

Finite Element Method Magnetics simulation and magnetic field measurement

To evaluate magnetic fields generated by coils, we use Finite Element Method Magnetics software (version 4.2) to simulate magnetic field of coil in 2D (Fig. S2B, S8C). In program setting, we used magnetics problem mode, planar configuration, and centimeter as unit in FEMM4.2. Copper wire (AWG12, AWG18) parameters and air condition were from the materials library in FEMM4.2. Gauss meter (TM801, KANETEC) was used for magnetic field measurement in the coil. The axial probe was used to detect magnetic field that generated inside of coil.

Cell viability test

The death of primary cultured neurons with multiple magnetomechanical stimulations were measured with propidium Iodide (PI; P1304MP, Invitrogen). After imaging the $Ca^{2+}$ responses of primary cultured neurons with Fluo-4, PI was added to the extracellular solution to achieve a final concentration at 120 μM. After 5 min, the fluorescence of Fluo-4 and PI were imaged by using fluorescence microscope (SS-1000-00, Scientifica). Thorlabs light source (LEDD1B), Hamamatus C13440 camera, and filter cubes (39002 and 39010, Chroma) was used.

Magnetic apparatus for in vivo experiments

To generate a uniform magnetic field for wireless neuronal stimulation *in vivo*. Four custom-made air-core coils were used for in vivo experiments. Each coil has 500 turns of 12 AWG cooper wire. The resistance and inductance of each coil is 1.02 to 1.55 Ω and 22 to 31.6 mH, respectively. The inner-diameter, outer-diameter, height of each coil is 20 cm, 28 cm and 5 cm, respectively (Fig. S8A). As demonstrated in Figure S8C,D, four coils are separated into two pairs which stack on each other. The gap between coil at center is 4cm. Full bridge modules (AQMH3615NS, AKELC) was used as the drivers for each coil. The drivers were controlled by the 5 V square waves generated by Arduino UNO (Arduino). The Arduino were controlled by custom-made script with

Arduino IDE. Internal power supply is 5V and provide by Arduino to give working voltage for full bridge modules. External power supply (HJS-1000 model, HuntKey) was used for supplying the currents for the coils. 10 A currents were used for generating a uniform magnetic field with 50 mT at 10 Hz in the coils for *in vivo* experiments. The temperature change less than 1 ℃ in 2 minutes with 10A DC current (Fig S8E).

Statistical analysis

All statistics were performed in JASP (v0.14.1.0, JASP team). All the error bars in the bar graphs indicate standard error of mean (s.e.m.). All the gray or pink area in the traces of fluorescence changes indicate standard error of mean (s.e.m.). Wilcoxon signed-rank test was used for comparing paired data in Figure 1J, 4E-F, and 4H-I. Tukey post-hoc test and Two-way ANOVA was used for comparing data with two factors in Figure 2E. Dunn post-hoc test and Kruskal-Wallis test were used for comparing data in Figure 2J-K and 3E-F. Wilcoxon rank-sum test was used for comparing unpaired data in Figure 4G and 4J.

Arduino code for controlling in vivo coil system

Arduino Uno board was used to control the alterative magnetic field for *in vivo* coil system. The code was written by Arduino IDE (version 1.8.13) and was compiled for Arduino Uno and Arduino Mega. In this program, phases of two signals for 4 full-bridge drivers had 180° difference. The pin 4 to 7 and of Arduino Uno were used for one signal to generate same orientation of magnetic fields in 4 coils. The pin 8 to 11 of Arduino Uno were used for the other signal to generate the reversed magnetic fields in 4 coils. Serial BUAD rate set at 500000 for stimulation trigger and monitor on computer.

```
void setup() {
  Serial.begin(500000);
  pinMode(4,OUTPUT);//control1
  pinMode(5,OUTPUT);//control2
  pinMode(6,OUTPUT);//control3
  pinMode(7,OUTPUT);//control4
  pinMode(8,OUTPUT);//control1
  pinMode(9,OUTPUT);//control2
  pinMode(10,OUTPUT);//control3
  pinMode(11,OUTPUT);//control4
  pinMode(13,OUTPUT);//indicate light
}
void loop() {
  // put your main code here, to run repeatedly:
  allStop();
  readFromPC();
}
void freqstart(){
  digitalWrite(4,HIGH);//control1
  digitalWrite(5,HIGH);//control2
  digitalWrite(6,HIGH);//control3
  digitalWrite(7,HIGH);//control4
```

```
  digitalWrite(8,LOW);//control1
  digitalWrite(9,LOW);//control2
  digitalWrite(10,LOW);//control3
  digitalWrite(11,LOW);//control4
  digitalWrite(13,HIGH);//test
  delay(100);//100->5Hz; 50 -> 10hz; 25 -> 20hz
  digitalWrite(4,LOW);//control1
  digitalWrite(5,LOW);//control2
  digitalWrite(6,LOW);//control3
  digitalWrite(7,LOW);//control4
  digitalWrite(8,HIGH);//control1
  digitalWrite(9,HIGH);//control2
  digitalWrite(10,HIGH);//control3
  digitalWrite(11,HIGH);//control4
  digitalWrite(13,LOW);//test
  delay(100);//100->5Hz; 50 -> 10hz; 25 -> 20hz
}
 void allStop(){ // Both IN1 & IN2 LOW
  digitalWrite(8,LOW);//IN2
  digitalWrite(9,LOW);//IN2
  digitalWrite(10,LOW);//IN2
  digitalWrite(11,LOW);//IN2
  digitalWrite(4,LOW);//IN1
  digitalWrite(5,LOW);//IN1
  digitalWrite(6,LOW);//IN1
  digitalWrite(7,LOW);//IN1
}
void readFromPC(){
    String s = "";
    int i = 0;
    int j = 0;
    int k = 0;
    while (Serial.available()) {
        char c = Serial.read();
        if(c!=""){
        Serial.println(c);
        }
        if(c!='\n'){
          if(c == 'a'){
            Serial.println("start stimulate");
            for(j=1;j<=5;j++){
              Serial.print("doing ");
              Serial.println(j);
              for(i=0;i<150;i++){   //5hz=> i=150; 10hz=> 300 for 30s;
                freqstart();
              }
```

```
            Serial.print("done");
            Serial.println(j);
            allStop();
            delay(60000);
          }
          Serial.println("alldone");

        }

      }
      //delayMicroseconds(2);    //change BUAD to 500000, no need delay anymore
   }
}
```

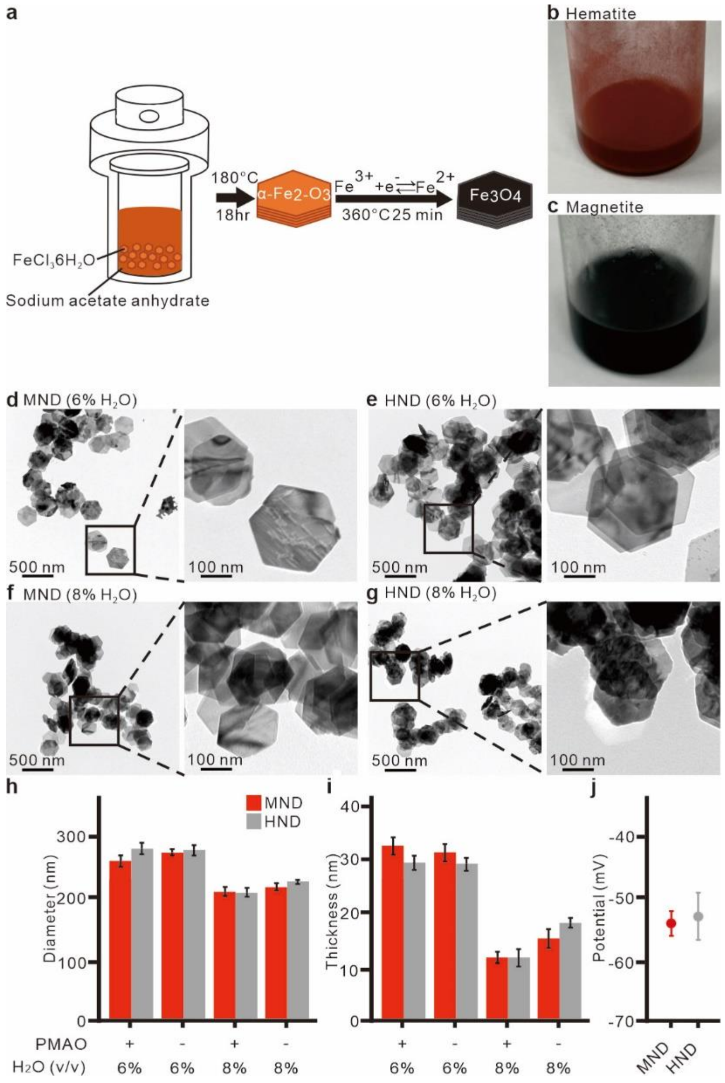


**Figure S1, Preparation of magnetic nanodiscs.**

**a**, Schematic of the two steps magnetic nanodisc synthesis process. **b**, A photo of HND in glassware from first step of reaction. **c**, A photo of MND in glassware from second step of reaction. **d**, TEM image of Magnetite nanodiscs with 6% $H_2O$ in the reaction solution at first step of reaction. **e**, TEM image of hematite nanodiscs with 6% $H_2O$ in the reaction solution. **f**, TEM image of Magnetite nanodiscs with 8% $H_2O$ in the reaction solution. **g**, TEM image of hematite nanodiscs with 8% $H_2O$ in the reaction solution. **h**, Diameter of nanodiscs measured from TEM images (n = 10). **i**, Thickness of nanodiscs measured from TEM images (n = 10). **j**, Zeta-potentials of PMAO-coated nanodiscs with 6% $H_2O$ in the reaction solution (MNDs, n = 12; HNDs, n = 9).

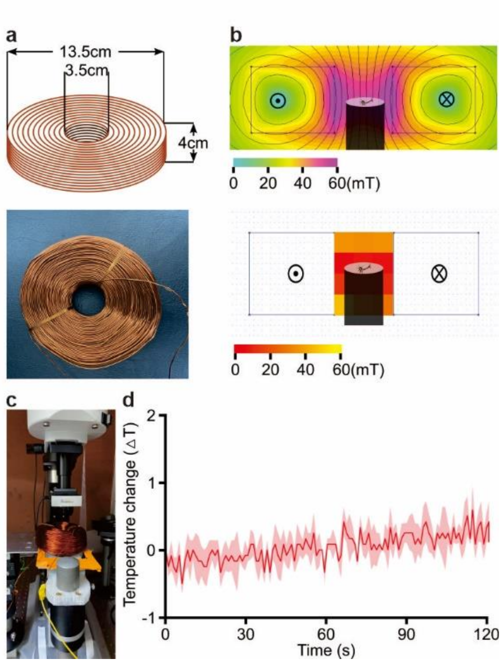

**Figure S2, Magnetic apparatus for magnetic field stimulation with upright microscope.**
**a**, Custom-made coil for upright microscope. Top, Design of the coil. Bottom, The photo of custom-made magnetic apparatus for *in vitro* fluorescence imaging. **b**, Heat maps of magnetic fields generated by coil. Top, FEMM simulation of the coil with 3A current. Bottom, the measured magnetic field inside of the coil with 3A DC current. **c**, The photo of custom-made coil set up at upright microscope. **d**, Temperature measured from the inner wall of *in vitro* coil.

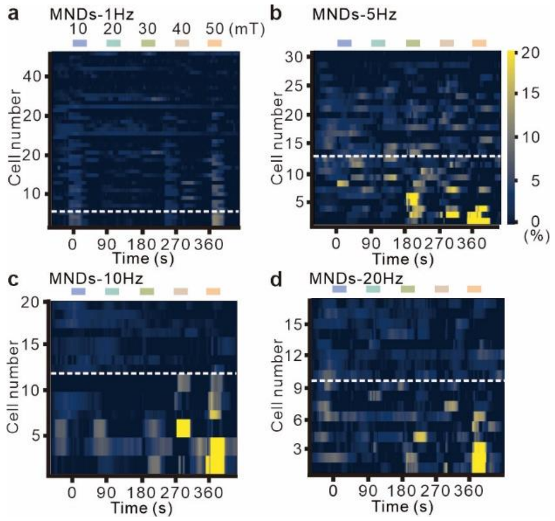


**Figure S3, The neuronal activity with different magnetic parameters.**
**a** to **d**, The heat maps of the calcium responses in individual neurons with magnetomechanical stimulation by using alternative magnetic field at di at 1 Hz (a), 5 Hz (b), 10 Hz (c), and 20 Hz (d). In each frequency, the magnetic field intensities were sequentially increased from 10 to 50 mT.

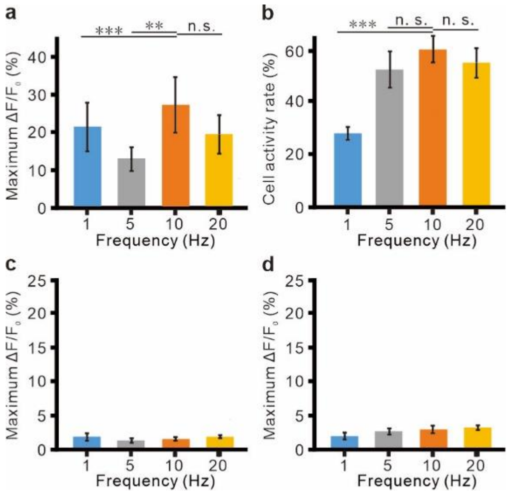


**Figure S4, The neuronal activity with different Frequency parameters.**

**a**, Bar graph of Maximum $\Delta F/F_0$ of MNDs treated neurons with 4 magnetic stimulation at different frequency. F = 1.05, p = 0.003, Kruskal-Wallis test. **p < 0.01, ***p < 0.001, Dunn post-hoc test **b**, Bar graph of cell activity rate of MNDs treated neurons with 4 magnetic stimulation at different frequency. F = 8.208, p = 0.005, Kruskal-Wallis test. ***p < 0.001, Dunn post-hoc test. **c**, Bar graph of Maximum $\Delta F/F_0$ of HND treated neurons with single magnetic stimulation at different frequency. F = 0.999, p = 0.268, Kruskal-Wallis test. **d**, Bar graph of Maximum $\Delta F/F_0$ of HND treated neurons with 4 magnetic stimulation at different frequency. F = 0.597, p = 0.392, Kruskal-Wallis test.

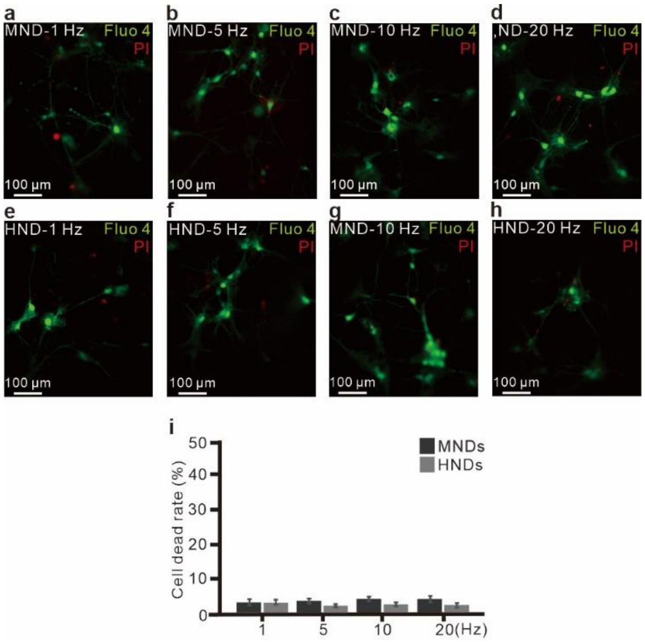


**Figure S5, Live-dead assay after Magnetomechanical stimulation.**
**a** to **d,** PI treatments in neurons with MNDs after magnetomechanical stimulation at 1 Hz (a), 5 Hz (b), 10 Hz (c) and 20 Hz (d). Green, Fluo-4; Red, PI. **e** to **h**, PI treatments in neurons with HNDs after magnetomechanical stimulation at 1 Hz (e), 5 Hz (f), 10 Hz (g) and 20 Hz (h). **i**, Quantify of live-dead assay with different frequency. Green, Fluo-4; Red, PI. $F = 0.003$, $p = 0.995$ for frequency; $F = 0.003$, $p = 0.862$ for type of nanodiscs; $F = 0.022$, $p = 0.995$ for interaction between frequency and nanodiscs; Two-way ANOVA.

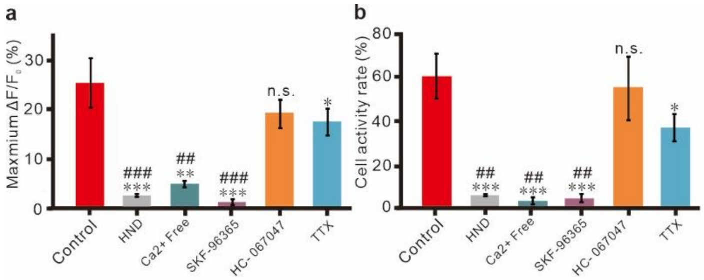


**Figure S6, Pharmacological dissection of Magnetomechanical stimulated response.**
**a**, Bar graph of Maximum $\Delta F/F_0$ of nanodiscs treated neurons with four magnetic stimulations. Except the HND group, MNDs are used in every other condition. $F = 7.767$, $p < 0.001$, Kruskal-Wallis test. *$p < 0.05$, **$p < 0.01$, ***$p < 0.001$, comparing to control group; ##$p < 0.01$, ###$p < 0.001$, comparing to HC-67047 group; Dunn post-hoc test. **b**, Bar graph of cell activity rate of nanodiscs treated neurons with four magnetic stimulations. Except the HND group, MNDs are used in every other condition. $F = 43.791$, $p < 0.001$, Kruskal-Wallis test. *$p < 0.05$, ***$p < 0.001$, comparing to control group; ##$p < 0.01$, comparing to HC-67047 group; Dunn post-hoc test.

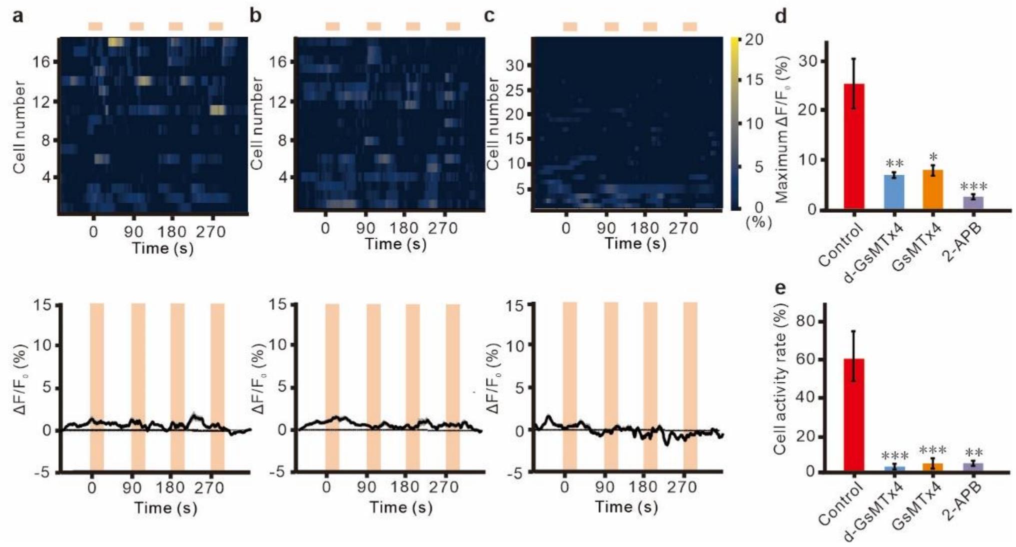


**Figure S7, Magnetomechanical stimulated response with GsMTx4, d- GsMTx4 and 2-APB.** **a** to **c,** The $Ca^{2+}$ responses by multiple magnetomechanical stimulation with bath application of GsMTx4 at 5 μM (a), d-GsMTx4 at 5 μM (b), 2-APB at 100 μM (c). The varying magnetic fields are 50 mT at 10 Hz for 30 s. Inter-stimulation intervals are 60 s. Top, heatmap of individual cells responses. Bottom, averaged fluorescence changes. The light orange areas indicate the time periods of magnetic stimulation. **d**, Bar graph of Maximum $\Delta F/F_0$ of MNDs treated neurons with different blockers. Control is same as Figure S6A. $F = 6.51$, $p < 0.001$, Kruskal-Wallis test. *$p < 0.05$, **$p < 0.01$, ***$p < 0.001$; Dunn post-hoc test. **e**, Bar graph of cell activity rat of MNDs treated neurons with different blockers. Control is same as Figure S6A. $F = 33.176$, $p < 0.005$ Kruskal-Wallis test. *$p < 0.05$, **$p < 0.01$, ***$p < 0.001$; Dunn post-hoc test.

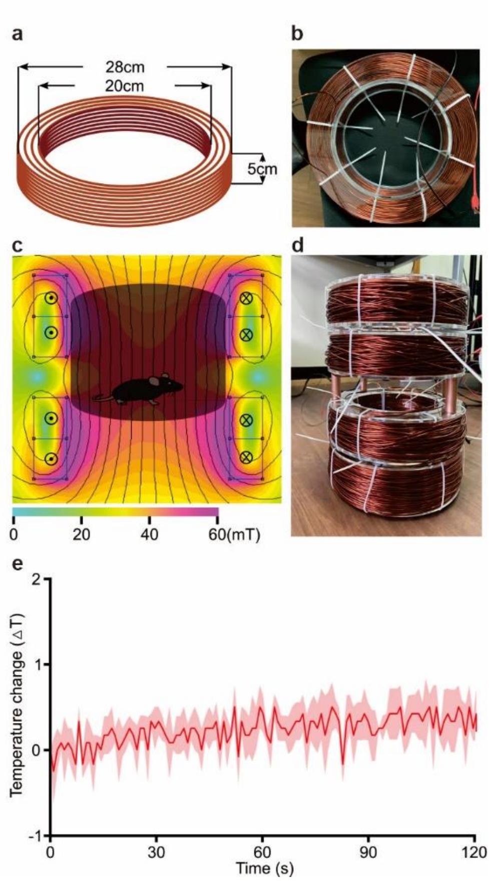


**Figure S8, In vivo coil system for animal test.**

**a**, Schematic of the design of single coil for *in vivo* experiment. **b**, The photo of a coil for custom-made magnetic apparatus for *in vivo* animal experiments. **c**, FEMM simulation of the in vivo magnetic apparatus with 10A current. **d**, The photo of custom-made magnetic apparatus for in vivo animal experiments. **e**, Temperature measured at the wall of *in vivo* coil with 10A DC current.

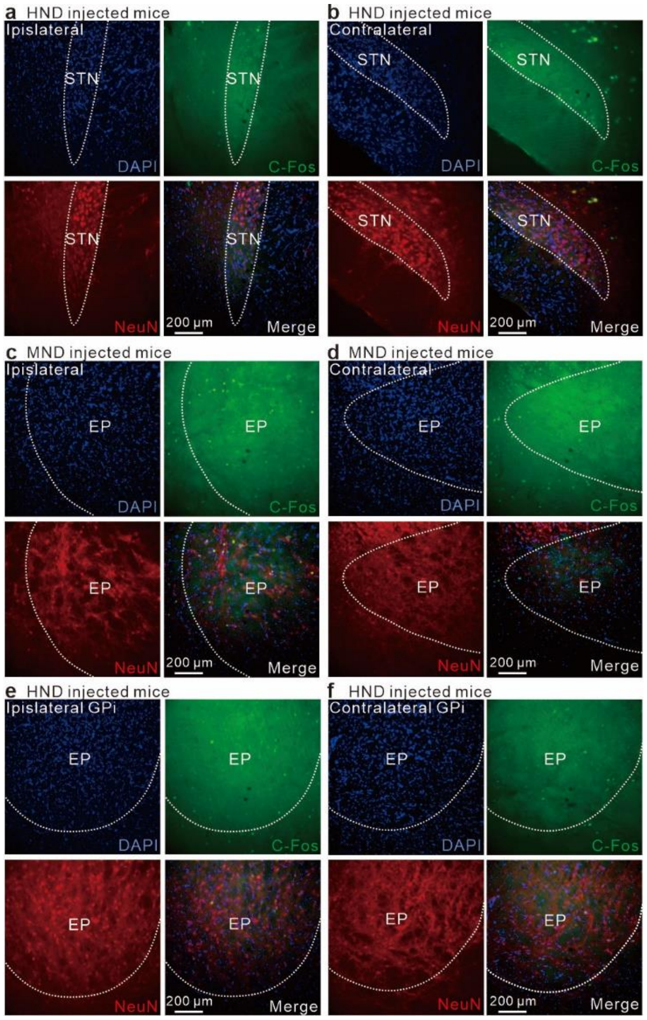


**Figure S9, c-fos expression in STN and EP of nanodiscs injected mice.**

**a** to **b,** Immunostaining of c-fos (top-left), NeuN (top-right), DAPI (bottom-left) and merged image (bottom-right) in STN after magnetic stimulations. Ipsilateral STN of HND injected mice (a). Contralateral STN of HND injected mice (b). **c** to **d,** Immunostaining of c-fos (top-left), NeuN (top-right), DAPI (bottom-left) and merged image (bottom-right) in EP after magnetic stimulations. Ipsilateral EP of MND injected mice (c). Contralateral EP of MND injected mice (d). **e** to **f,** Immunostaining of c-fos (top-left), NeuN (top-right), DAPI (bottom-left) and merged image (bottom-right) in EP after magnetic stimulations. Ipsilateral EP of HND injected mice (e). Contralateral EP of HND injected mice (f).
